# A high-throughput cost-efficient *in vitro* platform for the screening of immune senomodulators

**DOI:** 10.64898/2026.03.11.711033

**Authors:** Caterina Carraro, Timo Zajac, Sophie Lindenberg, Jacqueline Leidner, Alice Ragogna, Bidour Hussein, Sophie Mueller, Martina van Uelft, Heidi Theis, Elena De Domenico, Marc Beyer, Monique M.B. Breteler, Joachim L. Schultze, Anna C. Aschenbrenner, Jonas Schulte-Schrepping, Lorenzo Bonaguro

## Abstract

With advancing age, the immune system’s capacity to effectively combat pathogens diminishes. This decline of the overall immune function impacts both the innate and adaptive compartments, contributing in many cases to a systemic state of chronic inflammation which further increases the risk of the most prevalent non-communicable diseases and severe infections. Given the increase in median life expectancy with a demographic development towards a larger number of elderly people, identifying interceptive strategies to mitigate the individual and societal impact of diseases related to immune aging is of paramount importance. We developed a molecularly defined strategy to guide interventions with the aim to reduce immune aging. We introduce an omics-based drug screening platform to identify and characterize the pharmacological profile of immune senomodulators applicable to cross-age human cohorts using human-derived peripheral immune cells. To this aim we developed a robust experimental approach to screen for effective anti-aging compounds directly on human cells. This methodology allows us to quickly screen for drug candidates at different scales: from cost-effective bulk transcriptomics for a broader high-throughput overview of cellular responses, down to single-cell resolution approaches for a more detailed look at gene expression and other molecular data. This in vitro screening method is designed to maximize the clinical relevance of our findings, providing a direct link between preclinical research and patient care. By analyzing how different compounds affect the immune cells of individual persons, we can identify treatments that are most likely to be effective against aging in a subject-specific manner—a key step toward personalized medicine. In short, our approach enables a faster translation of anti-aging immune treatments from the lab to the clinic, tailoring them to each individual’s unique biological makeup.

## Introduction

The worldwide prolongation of human life expectancy is accompanied by an increase of age-related diseases such as cancer, autoimmunity, type 2 diabetes or neurodegeneration ^1^. The immune system has been established as a critical gatekeeper against such age-related diseases. Different mechanisms of immune dysregulation have been reported to contribute to the development of the most prevalent non-communicable diseases (NCDs) such as cardiovascular diseases ^2^, chronic respiratory diseases ^3^, neurological disorders ^4^, cancer ^5^, type-2-diabetes (T2D) ^6^ or autoimmunity ^7^, strongly suggesting that the immune system itself should be a key target for preventative interventions.

Aging itself is related to a multifaceted decline in both innate and adaptive immune functions through a process characterized by several hallmark alterations ^8^. While to some extent physiological, various conditions—from genetic predisposition to environmental exposure—can accelerate immune aging, leading to a premature decline in immune function where pharmacological intervention might be particularly suited to maintain or re-establish immune function. One aspect of immune aging, *immunosenescence,* describes a gradual functional decline of the adaptive immune system, marked by i) a significant reduction in the thymic output of naïve T cells leading to diminished T cell repertoire diversity, ii) impaired T cell receptor (TCR) signalling resulting in suboptimal activation and effector functions, and iii) the progressive accumulation of senescent immune cells contributing to tissue dysfunction ^8–10^. Another aspect is *inflammaging*, which represents a systemic shift towards a chronic, low-grade pro-inflammatory state, characterized by elevated levels of pro-inflammatory markers even in the absence of obvious infection or other immune stimulation. This process further contributes to the decline of whole-body functions, impacting various physiological processes and increasing the susceptibility of the aging population to NCDs.

Omics technologies, particularly transcriptomics, can provide a powerful readout for early drug discovery and repositioning, given their ability to capture the heterogeneity of complex phenotypes such as immune aging ^11^.

In this study, we assessed the impact of selected *senomorphics* and *immunomodulatory* agents with known or yet undescribed anti-aging activity on both the adaptive and innate immune system using primary human biomaterial. We innovated a throughput- and dimensionality-spanning platform including three distinct transcriptomic approaches – based on bulk, multi-omics, and single-cell - on human-derived blood immune cells, to identify and validate drug candidates for preventing or decelerating immune aging.

Considering the continuously decreasing cost and increased standardization of omics assays, this work provides a crucial foundation for future personalized medicine. This will enable the use of individual profiles to guide anti-aging pharmacological interventions, tailoring treatments to unique aging trajectories and susceptibilities. Such an approach holds immense promise for accelerating the development of effective immune aging therapies and enhancing the overall well-being of our increasingly aging global population.

## Results

### Cost-optimized bulk transcriptomics identify drugs with potential anti-aging activity

To first identify compounds with anti-immune aging activity, we designed a *proof-of-concept* pool of n=8 potential immune aging modulators including approved or investigational drugs as well as an endogenous metabolite. The selected candidates were previously reported to either induce i) an aspecific anti-aging effect (senomorphics) or ii) an immunomodulatory effect without information on their potential senomodulatory activity on the peripheral immune compartment. Specifically, we selected rapamycin, spermidine, and metformin as *senomorphics*, and imiquimod, dexamethasone, celecoxib, methotrexate and losmapimod as known *immunomodulators* ^9,12–18^. All these compounds have been previously suggested to exert an effect on the immune system but with only partial description of their overall activity profile and mechanism of action in the context of counteracting immune aging. For example, spermidine, an endogenous metabolite, is currently undergoing clinical evaluation for its potential to counteract immune aging and enhance vaccination efficacy in elderly individuals ^19^. However, the precise mechanisms responsible for these beneficial effects seem to go beyond spermidine’s known capacity to induce autophagy and need to be further elucidated^20^. This selection was used to establish a cost-effective high-throughput protocol for investigating anti-aging activities based on bulk transcriptomics of primary human material. As depicted in Fig 1A, selected compounds were tested on PBMCs from n=4 healthy individuals at n=2 distinct 10-fold different concentrations (drug-specific and selected based on the existing literature) and n=3 timepoints (2 h, 6 h, 24 h), with the aim to capture the time dynamics of drug responses. At this broad screening stage, we included only young adult individuals (20-30 years old) to minimize inter-individual diversity within the immune system, known to increase with advanced age ^21^.

**Figure 1-.**
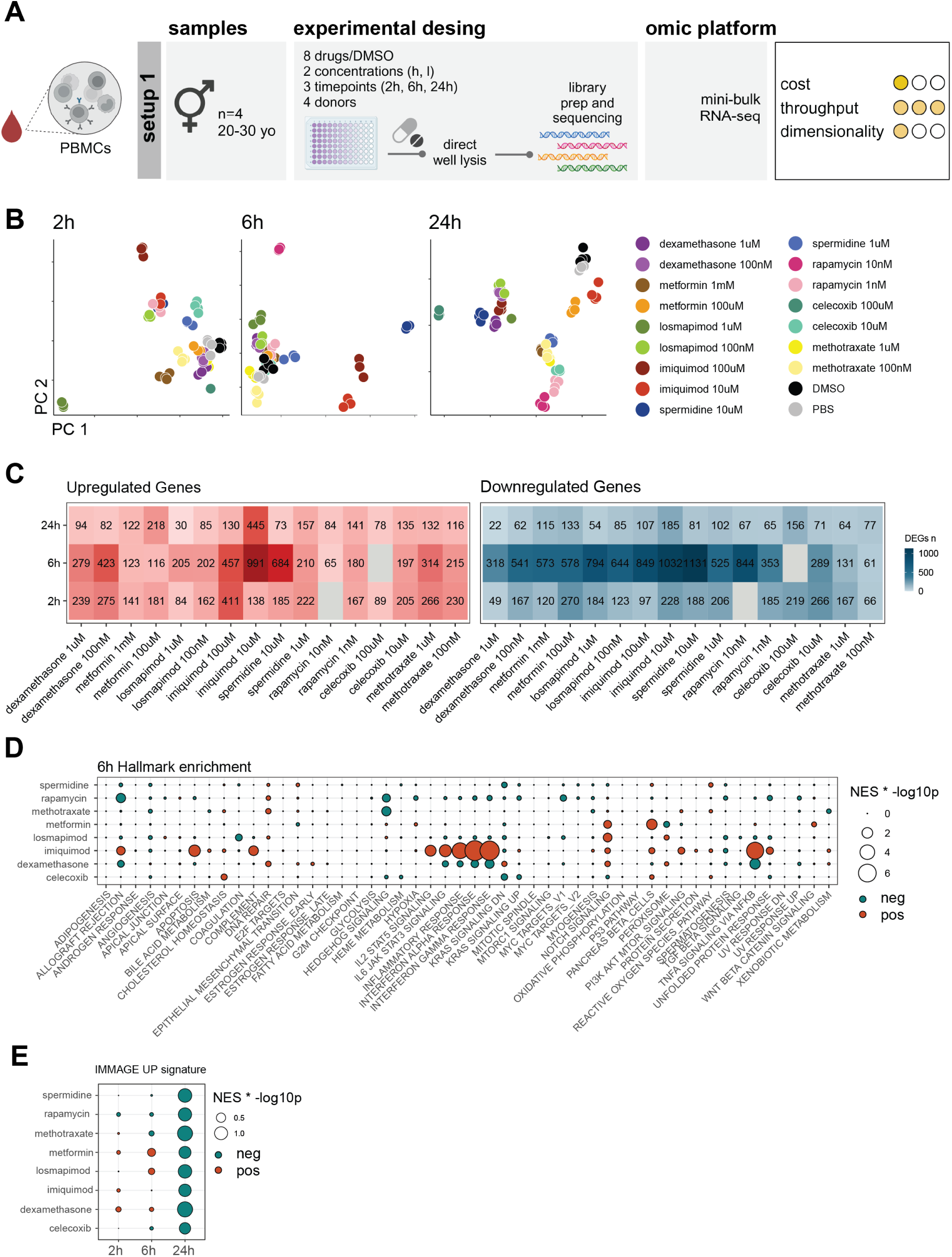
Mini-bulk RNA sequencing is a cost-effective approach to screen compounds for anti-aging effects. **A** Schematic representation of the first experimental design for the high-throughput low-budget drug screening on human PBMCs using a bulk RNA-seq readout (Study samples n=4, 2 males and 2 females). Created with BioRender. **B** PCA dimensionality reduction of the generated young human PBMCs dataset (colored by treatment and concentration tested, *granular annotation).* **C** Number of DEGs called for each treated versus ctrl comparison, drug treatments and time points (p.adj<0.05, FC_threshold_=2). **D** Enrichment analysis of the hallmark gene sets for each treatment after 6 hours of stimulation, dot size represents the product of the NES and the –log10 p value, dots are colored according on the regulation of the signature. **E** Enrichment analysis of the IMMAGE UP signature at 2, 6 and 24 hours of treatment. *PC: Principal Component; NES: Normalized Enrichment Score*

With this approach, we first sought to evaluate whether our bulk transcriptomics workflow was suitable to detect the anti-aging activity of selected drugs *in vitro*, thereby making the method applicable to transcriptomic screening of compound libraries at a reduced cost by optimizing for minimal cellular input and sequencing depth while maintaining sufficient information content. Second, we aimed at evaluating the anti-aging activity of test compounds on human peripheral blood mononuclear cells (PBMCs) as a function of treatment concentration and time point to further optimize downstream conditions for more fine-granular *in vitro* perturbation. Screening for n=4 individuals with n=8 compounds for n=3 time points at n=2 concentrations sums up to more than 200 individual bulk transcriptomes, requiring optimized protocols to maintain the affordability of the screening. The initial evaluation and quality control (QC) of the sequencing data highlighted that despite the intentionally reduced sequencing depth to 1 million reads per sample ^22^ most of the total 216 screened samples (184/216) passed the QC cutoff of half-million uniquely aligned reads per sample, defined as inclusion criterion for downstream analysis (Figure S1A), and, capturing biologically relevant protein-coding genes (Figure S1B). Following initial data normalization and transformation steps, we evaluated the overall effect of each drug perturbation on the transcriptome of PBMCs at three different time points, namely 2 h, 6 h and 24 h, to span across earlier and later transcriptional responses (Figure 1B). Principal component analysis (PCA) revealed treatment-specific transcriptomic changes, demonstrating dose- and time-dependent effects. As expected only a minimal variation was observed between DMSO and water controls, supporting the robustness of the perturbation obtained with the different treatments (Figure 1B). Differential expression analysis clearly revealed that most of the compounds showed the highest gene perturbations at 6 and 24 hours while Imiquimod and Dexamethasone were already active at 2 hours (Figure 1C, FC=2, p.adj<0.05). Over-Representation Analysis (ORA) for the *hallmark* gene sets showed regulation of immune relevant pathway after 6 h of treatment, including upregulation of inflammatory pathways in imiquimod-treated cells paired with a downregulation of the same pathways by dexamethasone (Figure 1D). We observed an overall increase in the regulation of several immunological pathways after 24 h (Figure S1C).

To understand the impact of the selected treatment on immune aging, we performed an enrichment analysis using a previously published bulk PBMCs gene signature (IMMAGE ^23^) recapitulating transcriptomic changes associated with peripheral immune aging. To note, all the tested compounds exhibited an anti-aging activity, as indicated by a reduction of the IMMAGE upregulated signature, with dexamethasone and methotrexate showing the strongest effects (Figure 1E). We did not observe any relevant modulation of the IMMAGE downregulated signature. Consistent with previous reports ^24^, this signature is likely driven by differences in cellular composition rather than cell-intrinsic molecular changes, a feature that cannot be effectively recapitulated when using a PBMCs-based *in vitro* model (Figure S1D). In summary, we demonstrated that our cost-effective protocol could capture the pharmacological and anti-aging activity of the tested drugs. Furthermore, this initial screening allowed us to identify both the most relevant time points and concentrations to achieve such effect across compounds.

### Systematic description of peripheral anti-aging drug activity across immune compartments using single-cell multi-omics

Bulk transcriptomics holds significant potential for high-throughput cost-effective screening of compounds, as demonstrated for anti-aging candidates. However, when investigating the anti-aging activity of test molecules in heterogeneous tissues such as whole blood, it is crucial to understand which specific cell types or cellular states are affected by the treatment. To obtain a fine-granular overview of drug activity profiles on the adaptive and innate peripheral immune compartments, we established a second complementary setup based on a single-cell multi-omics approach, thus increasing the dataset dimensionality while maintaining a relatively high throughput. For this experiment, we selected effective concentrations and time-points based on our initial screening (Figure 2A). Consequently, we tested the selected drugs at the defined time points on PBMCs from four donors within the same age range as in the bulk experiment (aged 20-30). We performed this experiment using the BD Rhapsody HT system, an advanced version of the original BD Rhapsody platform that allows for the simultaneous analysis of up to 96 samples in a single experiment. To also investigate the impact of the selected compounds on the surface proteome we combined the transcriptome analysis with epitome analysis using the BD ABseq protocol, leading to the collection of a highly comprehensive multi-omics dataset describing the effects of the selected drugs.

**Figure 2-.**
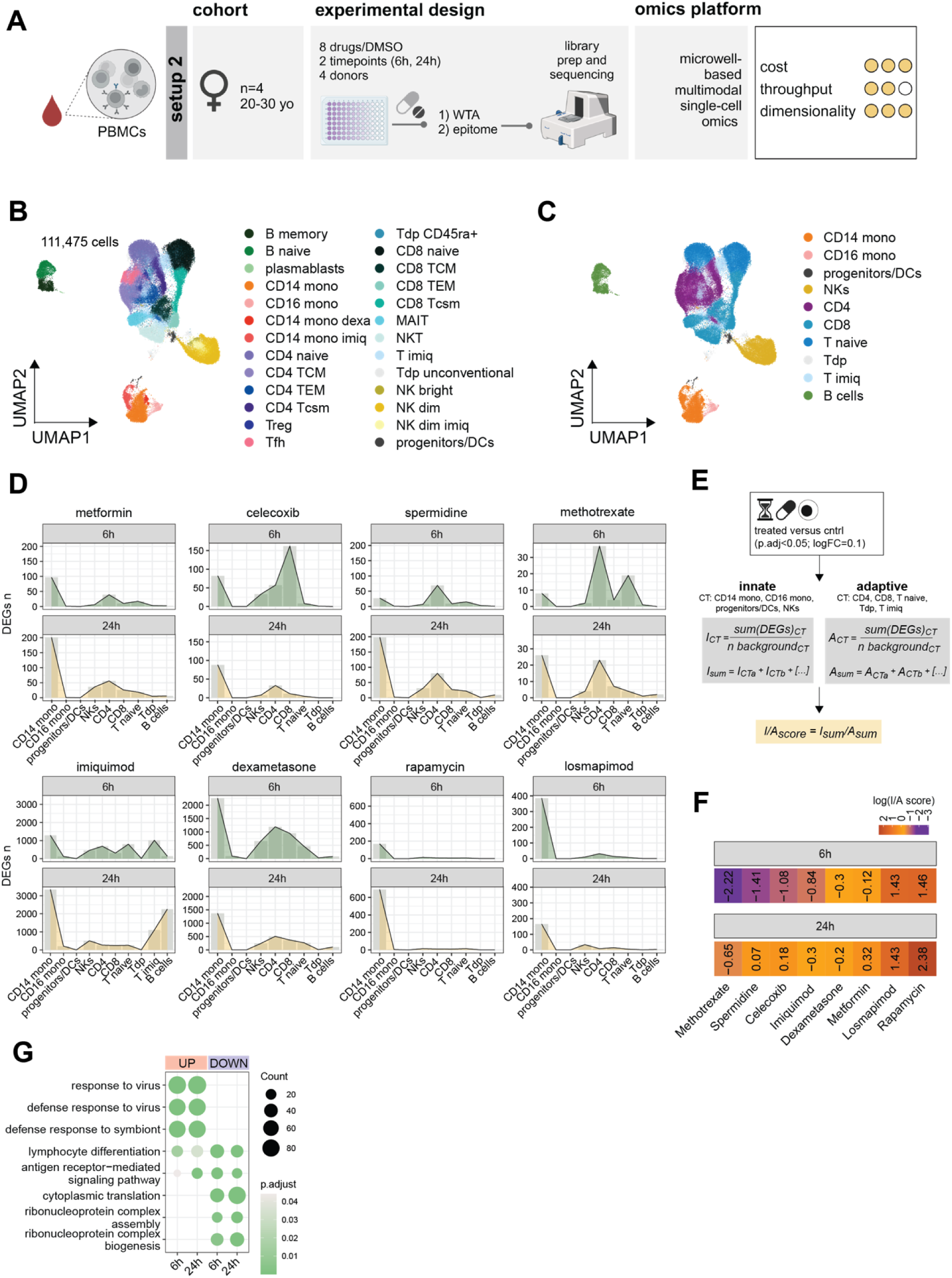
Single-Cell Transcriptomics Reveals Cell-Type-Specific Drug Effects. **A** Schematic representation of the second experimental design for the high-dimensional drug screening on human PBMCs using a single cell microwell-based multiomics readout. Created with BioRender. **B** UMAP representation of the generated young human PBMCs single-cell dataset (WTA and ABseq surface proteome integrated profiles, res=0.5, *granular annotation).* **C** UMAP representation of the generated young human PBMCs single-cell dataset (WTA and ABseq surface proteome integrated profiles, resolution=0.5, *coarse annotation*). **D** Time-specific drug effect dynamics reported as total number of detected DEGs (p.adj<0.05; logFC_threshold_=0.1) upon in vitro perturbation (6h or 24h) across the main PBMCs cell types. **E** Schematic representation of the strategy used to generate I/A scores: background-corrected DEG numbers are calculated per cell type and summed up separately for the innate and adaptive compartments. The ratio between the former and the latter is then generated per treatment and time point separately, enabling a relative comparison between tested compounds and timepoints. **F** Time-specific I/A scores describing the preferential activity of each tested drug on either the innate or the adaptive compartments. **G** ORA for the sum of up- and downregulated DEGs after treatment with imiquimod for the *T imiq* cluster (GO biological process, p.adj<0.05). **H** Schematic representation of the strategy employed to select the input genes used for the hdWGCNA analysis: per each drug and cell type, the top100[abs(logFC)*-log10(p.adj)] DEGs (separated for logFC>0 or logFC<0) were selected and summed up separately for the innate and adaptive compartments. *WTA: whole transcriptome analysis; ABseq: surface proteome analysis; mono: monocytes; dexa: dexamethasone; imiq: imiquimod; TCM: central memory T cells; TEM: effector memory T cells; Tscm: T memory stem cells; Treg: regulatory T cells; Tfh: follicular helper T cells; Tdp: double-positive T cells; DEG: differentially expressed gene; CT: cell type; I: innate; A: adaptive; ORA: over-representation analysis; GO: gene ontology; hdWGCNA: high-dimensional weighted correlation network analysis*.

We initially evaluated the quality of the sequencing libraries, which was consistent with the expected performance of the BD Rhapsody workflow (Figure S2A). Of note, a total of 111,475 high quality cells were recovered. The final dataset exhibited remarkably consistent quality control metrics across the different treatments and time points (on average 2,869 Unique Molecular Identifier (UMI) counts per cell and approximately 1,494 distinct genes per cell) (Figure S2A). Uniform Manifold Approximation and Projection (UMAP) dimensionality reduction revealed a substantial overlap between the different treatment groups, with the notable exception of Imiquimod, where a distinct shift in the T cell compartment was observed (Figure S2B). Based on the clustering results and the proteomic information of surface marker expression, we annotated all the major cell types expected in human PBMCs (Figure 2B-C, S2C) at two levels of granularity (Figure 2B-C). This analysis revealed that imiquimod induces substantial changes in the transcriptome of PBMCs, giving rise to specific clusters of T cells (*T imiq*) and classical monocytes (*CD14 mono imiq*). Similarly, we identified a cluster of monocytes responding to dexamethasone treatment (*CD14 mono dexa*). The presence of drug-specific clusters was further confirmed by frequency analysis (Figure S2D).

We next performed a differential gene expression analysis to evaluate the cell type-specific effects of each treatment (logFC=0.1, p.adjust<0.05). Each treatment resulted in a substantial number of differentially expressed genes, particularly following dexamethasone and imiquimod treatment (Figure 2D). Interestingly, we detected an overall higher number of differentially expressed genes at the 6 h time point compared to 24 h, highlighting a strong early response to the treatments, consistent with our previous findings in the bulk-based setup experiment (Figure 1). We also identified distinct activity dynamics on the innate versus adaptive immune compartments at the two time points across treatments. As an example, celecoxib and methotrexate exhibited a stronger effect on adaptive immune cells at 6 hours, while preserving a similar baseline activity on the innate compartment at 24 hours. To summarize this relative preferential activity on the innate versus the adaptive compartments, we generated an Innate/Adaptive (I/A) score, calculated as the ratio of background-corrected DEGs on innate versus adaptive cells (Figure 2E). Using this score, we were able to recapitulate the differential activity of each drug across different time points, further emphasizing the importance of incorporating this layer of cell type-specific activity analysis for the repurposing of pharmaceuticals in complex tissues like the peripheral immune system (Figure 2F). Overall, the I/A scores confirmed a general stronger response of the adaptive compartment at 6h, then shifting towards the innate compartment over time, with methotrexate being particularly active on the adaptive compartment and rapamycin on innate immune cells. Due to the previously identified dominant effects of imiquimod on the T cell compartment (Figure S2B), we evaluated the most affected pathways via ORA also in the T cells in the single-cell transcriptomics data. Being a TLR7 agonist, imiquimod potently induced an interferon I response, with the upregulation of terms involved in the response to viruses and T cell differentiation (Figure 2G). Considering the disruptive effect of imiquimod even at lower concentrations, specifically impacting the T cell compartment, we considered this treatment unsuitable for use in the aging population and excluded it from downstream analyses. Collectively, our findings reveal cell type-specific effects of the tested treatments, underscoring the critical importance of single-cell resolution when investigating potential new treatments.

### Using Single-Cell Multi-Omics to characterize and quantify drug mechanisms of immune senomodulation

To mechanistically characterize the effects of the treatments, with a specific focus on identifying both common and unique aspects of each, we performed co-expression network analysis (Figure 3A). For network construction, we selected the top 100 differentially regulated genes (upregulated or downregulated) in each cell type. Genes were ranked using the absolute product of the –log10 p-value and fold change ^25^, a metric that considers both the magnitude of change and statistical significance. Two distinct co-expression networks were generated: one associated with the innate immune compartment (3,091 genes), comprising eight modules, and the other associated with the adaptive immune compartment (2,933 genes), comprising seven modules (Figure S3A).

**Figure 3-.**
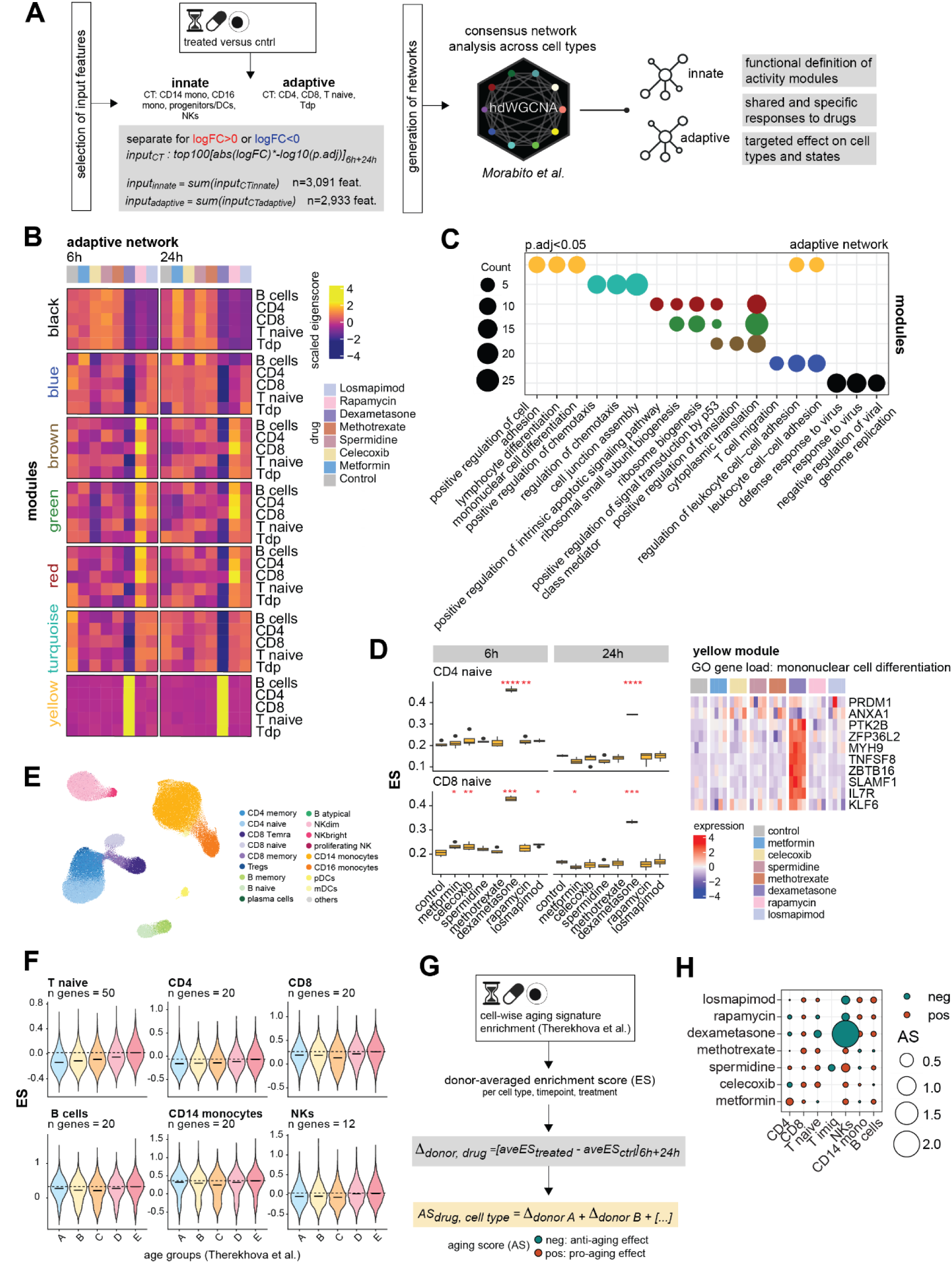
Co-expression Network Analysis Reveals Shared and Distinct Treatment and Drug Effects. **A** Schematic representation of the workflow applied to generate the *innate* and *adaptive* co-expression networks. **B** Heatmap reporting scaled eigenscores across drug treatments and time points for each of the gene co-expression modules identified within the *adaptive* network. **C** ORA across the gene modules identified within the *adaptive* network (GO biological process, p.adj<0.05). **D** Functional focus on the yellow *adaptive* module. Left: Donor-averaged ESs for genes included in the yellow *adaptive* module across conditions. Right: Heatmap reporting the expression of genes from the GO term load *mononuclear cell differentiation*. **E** UMAP representations of the Terekhova et al. PBMCs single-cell dataset. **F** ESs for the generated *senobase immune aging signatures across age groups in the* Terekhova et al. dataset. **G** Schematic representation of the strategy employed to generate ASs: donor-averaged enrichment scores are generated across cell types and states using previously derived immune aging signatures (*senobase*, ^43^), respective deltas are derived between treatment and control, summed up across time points and averaged across donors. **H** Summary ASs describing the anti- or pro-aging effect of different drug treatments on distinct immune cell types and states. *ORA: over-representation analysis; GO: gene ontology; ES: enrichment score; AS: aging score*.

We then investigated these networks in detail by characterizing the genes associated with each co-expression module. Within the adaptive cell network, several modules showed similar regulation across different treatments (Figure 3 B, C). Specifically, the blue, black, and turquoise modules were enriched for genes involved in adaptive immune system activation, including chemotaxis (turquoise, downregulated by multiple compounds at 6 h, particularly by dexamethasone), T cell activation (blue), and anti-viral responses (black, shared downregulated pattern between dexamethasone, metformin and losmapimod). In contrast, the brown, green, and red modules were preferentially regulated by rapamycin treatment, and the yellow module was almost exclusively regulated by dexamethasone treatment. Interestingly, the modules preferentially modulated by rapamycin showed similar enrichment for terms related to translation and ribosome biology, consistent with rapamycin’s role as a mTOR inhibitor. Most notably, the dexamethasone-upregulated yellow module was enriched for terms related to T cell activation and cell adhesion pathways, attesting its immunoregulatory, rather than purely immunosuppressive effect, in a context- and time-dependent manner ^26^. A detailed analysis of the yellow module shows a specific upregulation of this module after dexamethasone treatment in both CD4 and CD8 cells at 6 h and 24 h (Figure 3D). This module contained relevant genes for T cells homeostasis such as IL7R (CD127), ANXA1 (Annexin 1) and KLF6 (Figure 3D).

Analysis of the innate immune network (Figure S3B and S3C) also revealed both common and treatment-specific gene regulation. The black, blue, green, pink, and turquoise modules showed similar regulation across several treatments. Genes within these modules have important functions in coordinating the adaptive immune response, including regulation of mitochondrial biogenesis (black), response to bacteria or pathogen-associated molecular patterns (PAMPs) (blue), response to viruses (green and pink), and regulation of antigen presentation (turquoise), a key process for priming the adaptive immune response.

Modules with preferential regulation in only a few treatments were also identified. The brown module, preferentially regulated by rapamycin, was enriched for genes involved in translation and ribosome biogenesis ^27^. The red one, regulated by dexamethasone, included genes involved in metabolic regulation and steroid response. The yellow module, commonly regulated in response to multiple treatments and particularly after exposure of CD14 monocytes to losmapimod, showed enrichment for genes involved in the regulation of oxidative stress ^28^. As a validation step, since we used known compounds, our approach swiftly recapitulated the molecular activity of each of the tested compounds as described in the literature arguing that we captured ground truth for the effects of the compounds in our screening system. As a step forward, we focused on specifically evaluating the anti-aging activity of the tested drugs across different cell types. The IMMUN-AGE signature previously introduced was derived from a bulk analysis, thus not ideal to investigate cell type-specific anti-aging activities. We therefore extracted novel cell type-specific PBMCs signatures from a recently published single-cell study including 166 participants across the entire lifespan ^29^. First, we re-annotated the published dataset according to major cell types and lineages (Figure 3E). According to this annotation, we contrasted the group of older individuals (group E, aged 65 to 85) against all other younger age groups (A – aged 25 to 34, B – aged 35 to 44, C – aged 45 to 54, D – aged 55 to 64) and derived upregulated signatures of the basal transcriptional dynamics of immune aging (referred to as *senobase* signatures, see methods for details). We validated these signatures in the Terekhova dataset ^29^ showing a gradual increase in the signature enrichment with age (Figure 3F, S3D). Notably, the difference across age groups was more pronounced in long-living cells such as T cells and less strong in short-living cells such as monocytes and NK cells, as previously reported ^8^. Investigating the individual signatures, (Figure S3D) we identified previously reported genes linked to immune aging such as BCL2, HLA-DR, CST7 further supporting their biological relevance in aging processes.

We then evaluated the potential anti-aging effect of our treatments by investigating their ability to reverse the *senobase* signature in each cell type (Figure 3G; see Methods for details). This potential was assessed at both levels of cell annotation to monitor specificity of the observed effects on both, cell types and cell states (Figure 3H and S3E). Without vaccination, most of the tested compounds showed cell type-specific anti-aging effects, alternatively beneficial or unfavourable based on the considered cell type and state. Promisingly, dexamethasone showed an effect on naive T cells and NK cells, rapamycin on naive, NK and regulatory T cells, and spermidine was the only compound to show anti-aging activity in memory cells and regulatory T cells, all cell types whose function is particularly affected by aging. It is important to note, however, that we observed relatively high inter-individual variability in the anti-aging response to each drug (Figure S3F).

Collectively, our findings reveal the differential efficacy of the tested compounds in modulating immune aging across various cellular populations and states. This underscores the absolute necessity of including cell type-specific analyses in drug repurposing efforts, where single-cell multi-omics serves as a scalable approach to meet this goal.

## Discussion

Age-related diseases represent a global burden on healthcare systems which requires urgent attention as well as dedicated prevention and treatment programs ^33^. The aging immune system appears to be a key factor in defining the overall health state of an individual as well as in the aging process of other organs ^34^. Attempts to modulate or reverse immune aging include immune cell replenishment via adoptive transfer, antibody depletion, or modulation of the hematopoietic stem cell (HSC) niche ^34,35^. While all these approaches provided valuable insights into the mechanisms of immune aging and the potential for interventions targeting the immune system to prevent premature immune aging, their application in clinical practice remains limited.

At the same time, treatments with anti-aging compounds have been proposed, and some are in clinical trials ^36^. One example is metformin, originally conceived to treat diabetes, now being investigated for its potential to prolong the overall lifespan, with clinical trials examining its effects on cardiovascular function, immune function, and inflammation in older adults ^37^. Further compounds in clinical trials include nicotinamide mononucleotide (NMN) and nicotinamide riboside (NR), targeting the age-dependent decline of NAD+ level, crucial for cellular functions. The link between modulation of NAD+ and sustained physical performance, brain health and other age-related conditions is currently under evaluation ^38^. Additionally, rapamycin and related mTOR inhibitors, which showed anti-aging effects in different model organisms, are under investigation in clinical trials to assess their effect on immune function and metabolic health in the elderly ^12^. Finally, senolytic compounds, that target and eliminate senescent cells contributing to age-related decline, are undergoing clinical trials to assess their safety and efficacy in treating various age-related diseases.

To systemically extend these efforts to a broader range of chemical compounds and biologicals with potential immunomodulatory effects, more personalized pre-clinical screening approaches are needed. In addition, clinical trial–compliant assay systems with high granularity that generate high-dimensional data are required to prevent premature immune aging and optimize immunotherapies and vaccines in the elderly.

To address this gap, we developed a staggered approach combining different omics technologies (bulk, multi-omics, single cell transcriptomics), then utilized exemplary immunomodulatory compounds with potential efficacy on age-related immune deviation and illustrated the direct application in primary human biomaterial. This comprehensive approach allowed us to accomplish two significant goals: (i) to systematically characterize the immune-senomodulatory potential of various compounds, whether they act on general aging mechanisms or specific immune-related pathways, and (ii) to establish a scalable platform facilitating the design of more effective and personalized treatments for age-related immune decline.

The first level of our approach included the innovation of an optimized bulk transcriptomic protocol designed to reduce cost while capturing relevant biology which led to the identification of compound-specific transcriptional alterations (Figure 1). Here, we screened over 200 bulk transcriptomes derived from n=8 drugs, n=2 concentrations, and n=3 timepoints in n=4 donors allowing us to define optimal conditions for further fine-granular screening steps. This initial level would be easily scalable in a cost-effective manner to several thousand samples. We could further envision that AI-driven approaches to select compounds from larger databases or even from generative AI models ^39^ could be performed prior to this screening step to reduce the compounds to be tested in our system.

Optimal conditions for time and concentration were defined for each compound and applied on the next level of the approach, which is the application of single-cell multi-omics (single cell proteogenomics and whole transcriptomes), holding the potential to provide cell-type- and cell-state-specific information on compounds’ activity. Here, the major finding was that different cell types and states could exhibit both, accelerating and decelerating effects on immune aging processes, depending on the compound studied (Figures 2 and 3). Notably, dexamethasone, spermidine, and rapamycin emerged as promising drug repurposing candidates, demonstrating strong anti-immune aging effects in several cell types with only minor accelerating effects in others (Figure 3). Specifically, our single-cell analysis allowed us to dissect the heterogeneous responses of various cell populations to these compounds, providing a more nuanced understanding compared to traditional bulk approaches. Interestingly, we observed that dexamethasone and rapamycin potentially restored normal function in naive T cells, while spermidine’s activity was more pronounced in effector and memory cells, suggesting a potential for this treatment to reestablish existing immune memory ^31^ (Figure 3). The distinct cell-type specificities of these drugs suggest that combination therapies, or personalized treatments tailored to an individual’s immune profile, could maximize therapeutic benefits. Collectively, these findings highlight the power of single-cell resolution in identifying potential cell-specific toxicities or beneficial effects that might be masked by bulk data analysis.

Overall, our results support the idea of two possible complementary strategies for tackling this immune aging, each with a distinct focus and application. The first approach is a long-term, preventative strategy that aims to slow the natural decline of the immune system over a person’s lifetime. The second is a more immediate, temporary intervention designed to boost immune function in specific situations, such as before vaccination, to ensure a robust response. Our screening identified rapamycin and spermidine as suitable candidates for a preventive treatment, consistent with previous research. On the other hand, adjunctive treatment with dexamethasone shows potential for promoting a more coordinated immune response, as evidenced by an early increase in adaptive immune cell priming, pointing to a potential use of this compound as booster of vaccine response. These observations align with previous reports and are supported by our data ^40^.

Our study is limited by the initial selection of compounds, which was based on previously reported potential anti-immune aging activities. This work is intended to serve as a comprehensive blueprint for investigating much larger libraries of compounds. Furthermore, the sample size of elderly individual samples presented here was limited in number. Future studies with a larger sample size will have to follow investigating whether the observed variance is a function of distinct “aging endotypes”. Such aging endotypes might then be used for more effective, personalized therapeutic or preventative interventions.

Finally, modelling the response to vaccination using peripheral blood provides information only on the early phase of the response. While this early phase is critical for the correct orchestration of the later adaptive response and memory formation, the introduction of adaptive immune organoids ^41^ into this framework could provide valuable information on the late phase of the response as well. Although this addition would be extremely useful, current organoids are derived from human secondary lymphoid organs, which cannot be removed without medical indications. Future developments in the generation of such organoids, combining iPSC-derived cells and peripheral blood, could constitute a powerful tool for investigating vaccination responses in the elderly.

In summary, this study thoroughly investigated a specific selection of compounds with potential anti-immune aging activity. By employing a rigorous in-vitro drug screening method on human peripheral blood mononuclear cells (PBMCs), we effectively showed how transcriptomics could provide robust clinical indications of these compounds’ efficacy. Additionally, we established a multi-layered approach for drug discovery and repurposing that could serve as a blueprint for investigating other compounds and, more importantly, other conditions modellable in vitro.

## Methods

### Isolation and cryopreservation of PBMCs from human blood

Peripheral blood mononuclear cells (PBMCs) were isolated from EDTA-anticoagulated peripheral blood via density gradient centrifugation using Pancoll (density: 1.077 g/mL). The isolated cells were then washed twice with Dulbecco’s Phosphate-Buffered Saline (DPBS) and cryopreserved in RPMI 1640 supplemented with 40% fetal bovine serum (FBS) and 10% dimethyl sulfoxide (DMSO). Frozen PBMCs were rapidly thawed in a 37°C water bath, immediately diluted in pre-warmed RPMI 1640 supplemented with 10% FBS (Gibco) and centrifuged at 300 × g for 5 minutes. Following centrifugation, the cell pellet was resuspended in RPMI 1640 supplemented with 10% FBS and processed for downstream applications.

### In vitro treatment of human PBMCs

After thawing, cells were seeded onto 96-well plates at a density of 100,000 cells per well for the bulk transcriptomics experiment, a density typically sufficient to obtain robust RNA sequencing data from a population average. In contrast, a higher seeding density of 300,000 cells per well was employed for the single-cell experiment. This increased initial number is crucial because the subsequent detachment steps, necessary to obtain individual cell suspensions for single-cell capture and barcoding, inevitably lead to a significant loss of cells. Therefore, starting with a higher density ensures that a sufficient number of viable single cells are available for downstream processing and analysis. Immediately following treatment with the selected compounds (Table S1), the cells were stimulated and incubated at 37°C with 5% CO₂ for the desired duration, allowing the compounds to exert their effects before cellular RNA was harvested. Cells were maintained in RPMI medium supplemented with 10% FBS and 1% Pen Strep to provide the necessary nutrients and prevent microbial contamination. For longer incubation time points, cells were meticulously checked daily for any signs of bacterial contamination to ensure the integrity of the experiment. Both water control and DMSO control were included.

### Mini-bulk RNA sequencing - data generation

Following incubation for the defined time point (Table S2), 96-well plates were centrifuged at 300 × g for 5 minutes. The supernatant was carefully removed, and the cells were washed once with 200 µL of ice-cold PBS. The cell plate was then centrifuged again at 300 × g for 5 minutes at 4°C. After complete removal of the supernatant, cells were lysed with 15 µL of SMART lysis buffer (composition specified elsewhere) and incubated on ice for 5 minutes. The cell plate was then sealed, vortexed to ensure complete lysis, and centrifuged at 1,000 × g for 1 minute to collect all cell lysate. The cell plate was stored at −80°C until library preparation.

SMART-seq2 libraries were prepared according to Picelli et al. (2013) ^42^ and Leidner et al. (2024) ^22^, using a 6 µL volume of cell lysate as input. Library quality control was assessed using a TapeStation instrument. Libraries from individual wells were then equimolarly pooled, and the final concentration for sequencing was determined using a Qubit instrument (High-Sensitivity Assay - Thermo Fisher Scientific). Final libraries were loaded at a final concentration of 650 pM on a NovaSeq 2000 instrument using a P3 flow cell. Sequencing was performed paired-end with 51 base pairs for each read, including two 8-base pair index reads.

### Mini-bulk RNA sequencing - data pre-processing and analysis

Raw sequencing results were demultiplexed according to the index pairs using bcl2fastq (v. 2.20) with default setting. Sequencing reads were then aligned to the human genomes (Gencode Version 27) with Kallisto (v. 0.440) with default parameters. Downstream analysis was performed in R (v. 4.3.0) and within the DEseq2 analysis framework (v. 1.40.2). Other plots were generated using ggplot2 (v3.4.2). After filtering the samples with > 0.5 M pseudoaligned reads, samples were loaded in the DEseq2 workflow with the *tximport* functions separately for the three timepoints. After data loading the DEseq2 object was normalized and vst transformed. Experimental batch effect was removed with the indentification of surrogate variable with the SVA package (v. 3.48.0). The obtained surrogate variables were included in the DEseq2 model for differential expression analysis (pval 0.05, FC 0.5, IHW pval correction), for visualization the vst transformed count were corrected for the surrogate variables identified using the *limmaBatchEffectRemoval* function from the limma package (v. 3.56.2). Principal Component Analysis shown in Figure 1 was performed on all genes using the sva corrected counts as input. GSEA was performed with the *fgseaSample* function of the fgsea package (v. 1.26.0) using as gene rank the DEseq2 calculated fold change for each comparison and 10000 permutations for the calculation of the statistical significance. For visualization the product of the normalized enrichment score and the –log10 transformed p value were used. The IMMAGE signature was previously reported and curated ^23,24^, Hallmark enrichment was performed using the latest available signatures from the MsigDB database (20/05/2025), h.all.v2024.1.Hs.symbols.gmt. To ensure reproducibility of the analysis a containerized environment was used.

### Multi-omics single cell sequencing - data generation

Following incubation for the defined time point (Table S2), the 96-well plates were centrifuged at 300×g for 5 minutes. After carefully removing the supernatant, the cells were incubated for 15 minutes with 100 μL of EDTA (5mM in PBS) at 37∘C. The cells were then resuspended, and each well was rinsed with an additional 50 μL of EDTA buffer to minimize cell loss. Subsequently, the BD Rhapsody protocol was used to generate the final sequencing libraries, including whole transcriptome analysis, sample tags, and proteogenomics. For each BD Rhapsody lane, 12 samples were multiplexed to mitigate experimental batch effects. Final libraries were sequenced on an Illumina NovaSeq 6000 instrument using a NovaSeq S4 v1.5 flow cell at a final concentration of 1250pM. Sequencing was performed paired-end with read 1 at 51 base pairs and read 2 at 151 base pairs; additionally, two index reads of 8 base pairs were included.

### Single cell dataset pre-processing and downstream analysis

#### Newly generated single-cell datasets

For the multi-omics single cell dataset, raw sequencing data were demultiplexed according to the index pairs using bcl2fastq (v. 2.20) with default settings. Subsequently, sequencing reads were aligned to the human genome using the BD Rhapsody Pipeline (v.2.1) with default parameters, referencing the BD Bioscience annotation file RhapRef_Human_WTA_2023-02.tar.gz based on the Gencode v42 primary assembly.

Downstream analysis was conducted in R (v4.3.0) and within the Seurat single-cell analysis framework (v4.3.0.1). Other plots were generated using ggplot2 (v3.4.2). After filtering for high quality cells (% of mitochondrial reads<20, % of hemoglobin reads<5, % of ribosomal reads<25, number of genes>250 & number of UMIs>500), data were log-normalized and dimensionality reduction (PCA) was applied on most variable features, followed by clustering (resolution=0.5) and non-linear dimensionality reduction (UMAP). For the multiomics BD Rhapsody dataset, the ABseq and WTA layers were first batch-corrected for the treatment timepoint (Harmony v0.1.1) followed by generation of a joint embedding (*FindMultiModalNeighbors* function, dimensions 1:20). The ABseq information was primarily employed to annotate cell types and states in the multiomics BD Rhapsody dataset. Frequency analyses were performed by normalizing for the total number of cells. Differential expression analyses (DEA) were performed using the Seurat-embedded MAST function (p.adj<0.05, logFCthreshold=0.1, latent.vars=”donor_ID”). The *senovax* signatures were generated by comparing cells from the *old* versus *young* (in vitro) *vaccine-stimulated* groups, only considering female donors, to exclude the influence of sex-related genes only due to the absence of males in the *young* group. Over-representation analyses (ORA) were performed using the *compareCluster* function (clusterProfiler v4.8.1, p.adj<0.05, GO biological process). Enrichment scores per cell were calculated using the *AddModuleScore* Seurat function, while differences across groups were Wilcoxon-tested for statistical significance where reported (p<0.05, *stat_compare_means* by ggpubr v0.6.0). Heatmaps were generated using the ComplexHeatmap function (v2.16.0) after averaging the expression across all cells per donor, condition and cell type. I/A (innate/adaptive) scores were generated as summarized in Figure 2A: per each *treated versus control* comparison (drug-, timepoint- and cell type- specific), and separately for the innate and adaptive compartments, cell type-specific scores (ICT and ACT) were derived as the sum of DEGs (for that cell type across *treated versus control* comparisons) normalized on the number of genes expressed as background in the cell type of interest. Scores were then summed up per compartment, generating Isum and Asum values, whose ratio was then calculated per each drug and timepoint separately (I/A score). The network analysis was performed using hdWGCNA (v0.3.03), as summarized in Figure 3A and separated for the innate and adaptive compartments. Input genes for the network were selected as follows: per each *treated versus control* comparison (drug-, timepoint- and cell type- specific), and separately for logFC>0 or <0, the top100 genes (abs(logFC)*-log(p.adj)) were defined as summed up across timepoints. These discrete inputs were then summed up across comparisons of the innate versus adaptive groups, leading to a total of n=3,091 input genes (innate) and n=2,933 input genes (adaptive). Consensus network analysis was then performed using default parameters unless differently specified, metacells were generated (group.by = c(“anno_coarse”, “condition”), k = 20, max_shared = 10, ident.group = ‘condition’, min_cells = 45), soft power thresholds were then determined to construct both networks (minModuleSize=20) followed by calculation of eigengenes. AS (aging score) and VAS (vax aging score) were calculated as described in Figure 3G: cellwise enrichment scores were calculated using the respective cell type-specific Terekhova-derived *senobase* signatures, donor-averaged per donor, cell type/state, timepoint and treatment (aveES). Delta scores (Δdonor,drug) per generated per donor and drug by subtracting the respective untreated aveESctrl. Per drug and cell type/state, ASs were generated summing up the contribution of the different donor deltas (reported only if minimum 100 cells available per subgroup). The VASs across treatments were generated following the same principle but testing the *senovax* signatures on vaccine-exposed cells with drug prestimulation versus without (Figure S5 A).

#### Terekhova et al. dataset

For the generation of *senobase* signatures, the publicly available scRNA-seq dataset of Terekhova et al. was used. The dataset contains 1.9 million cells from 166 individuals aged between 25 and 85 years old. All participants were healthy and non-obese. We re-annotated the cell types in R (v. 4.3.0) using the Seurat single cell analysis package (v. 4.3.0.1). Annotation and subsequent analysis were conducted separately for each major cell population, including CD4 T cells, CD8 T cells, B cells, myeloid cells and NK cells. To minimized donor-driven effects, cell numbers for each individual were downsampled to the average cell number per donor. Data processing was performed using the Seurat pipeline beginning with normalization, variable feature selection, and scaling. TCR alpha and beta chains, as well as BCR light and heavy chains were excluded from variable features. The batch effect was corrected using Harmony (v0.1.1) based on the PCA results, and UMAP was computed on the Harmony-corrected dimensions. Cells were annotated by clustering using Seurat’s FindNeighbors and FindClusters functions (resolution: B cells = 0.1, CD4 T cells = 0.2, CD8 T cells = 0.1, NK cells = 0.5, myeloid cells = 0.2), followed by the identification of cluster marker genes. Based on the expression of these marker genes and known cell type markers clusters were assigned to their corresponding cell subtypes. For visualization purposes, 20% of cells from each cell type object were randomly subset, merged, and visualized as UMAP, following the workflow described above.

DEA was performed separately for B cells, CD14 monocytes, NK cells, naive T cells (CD4 and CD8 combined), CD4 T cells, and CD8 T cells (excluding naive cells). Individuals were grouped by age into ten-year intervals: A (25-34), B (35-44), C (45-54), D (55-64), and E (65+). All age groups were compared to the oldest age group (E) using Seurat’s MAST test (min.pct = 0.1, logfc.threshold = 0.1).

Based on the described DEA, *senobase UP* signatures were generated only considering DEGs from the comparisons A, B, and C versus E, separately for B cells, CD4, CD8, T naive, CD14 monocytes and NKs and excluding genes non expressed in target cell types of our dataset. We defined upregulated signatures of n=20 genes across cell types, n=50 for T naive cells due to their higher magnitude of deregulation. For NKs, due to the limited number of DEGs, we defined the signature as the sum of all upregulated DEGs (n=12). Hits were prioritized for their recurrence across multiple comparisons up to the definition of the target genes number: if more than 20 (T naive, n=50) upregulated DEGs were detected across the 3 comparisons (A, B, and C versus E), genes were further selected based on the lowest p.adj. In case of DEGs overlapping across more than one comparison, the hit with the lowerst *p.adj* was considered. If less than n=20 (T naive, n=50) DEGs were overlapping across the n=3 comparisons, the remaining genes were selected based on the lowest *p.adj*.

## Declarations

### Availability of data and materials

Raw sequencing reads from bulk transcriptomic and BD Rhapsody multi-omics generated during this study will be deposited at the European Genome-phenome Archive (EGA), which is hosted by the EBI and the CRG.

### Code availability

All the computational analyses were performed using the R programming language. Scripts will be uploaded on DZNE GitLab repository upon publication. Annotated Seurat object will be deposited on Zenodo upon publication.

### Authors’ contributions

Conceptualization was by L.B, J.S.S. and C.C. Donor recruitment and processing of biomaterial L.B, A.C.A and M.B. The methodology was devised by L.B, J.S.S., C.C., T.Z., S.M., M.v.U., E.D.D. and H.T. L.B., C.C., T.Z., S.L., J.L. performed formal analysis. L.B. and C.C. carried out the investigations. The draft manuscript was written by L.B, and C.C. All authors reviewed and edited the manuscript. Visualization was by C.C. and S.L. The project was supervised by L.B. Funding acquisition was by L.B, J.S.S, M.B, A.C.A and J.L.S.

---

**Figure S1-.**
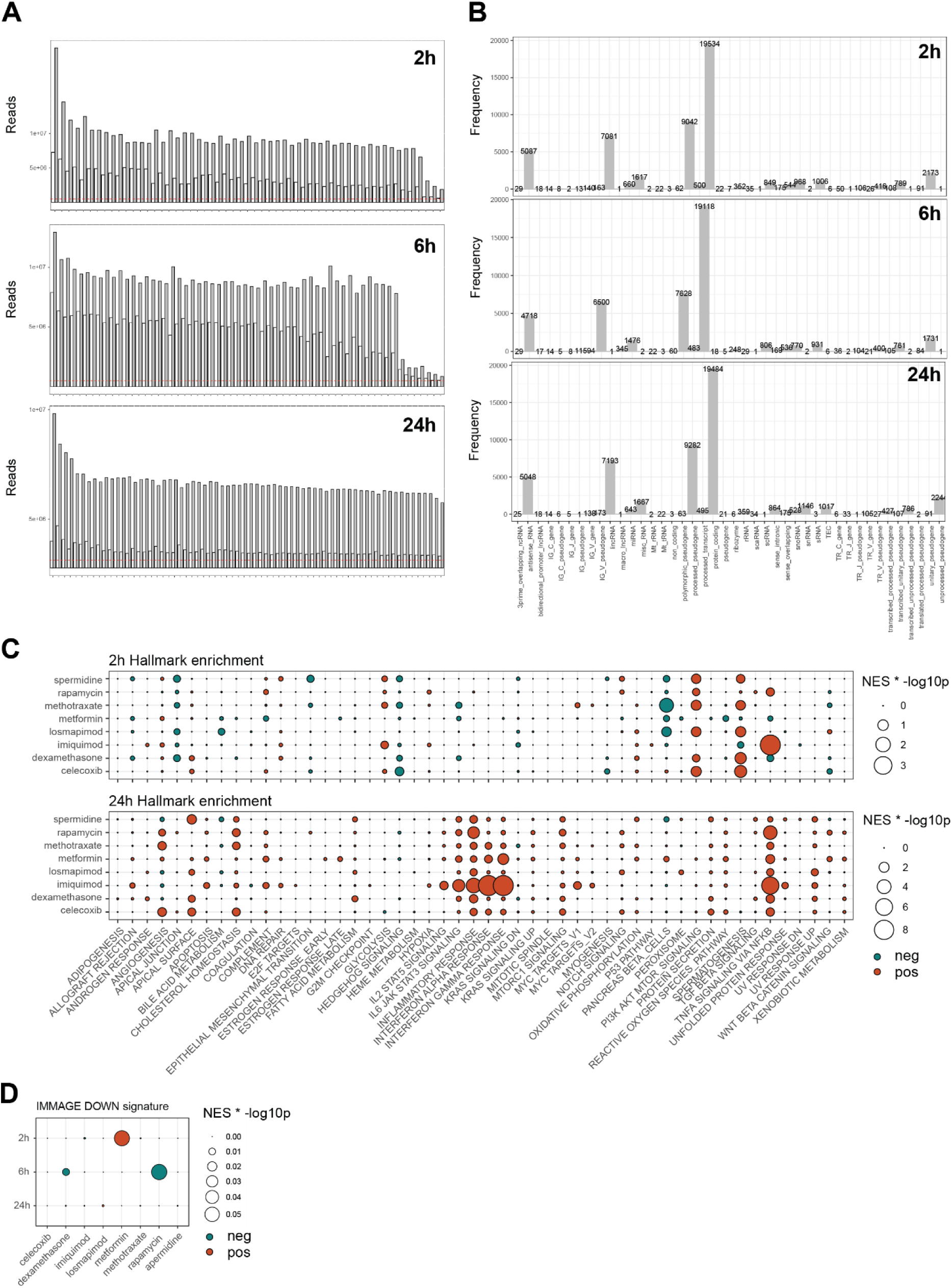
Mini-bulk RNA sequencing is a cost-effective approach to screen compounds for anti-aging effects. **A** Number of sequenced reads and pseudoaligned reads for each sample at 2, 6 and 24 hours after treatment. **B** Histogram of the number of detected genes splitted by the type of gene. **C** Enrichment analysis of the hallmark gene sets for each treatment after 2 and 24 hours of stimulation, dot size represents the product of the NES and the –log10 p value, dots are colored according on the regulation of the signature. **D** Enrichment analysis of the IMMAGE DOWN signature at 2, 6 and 24 hours of treatment. *PC: Principal Component; NES: Normalized Enrichment Score*

**Figure S2-.**
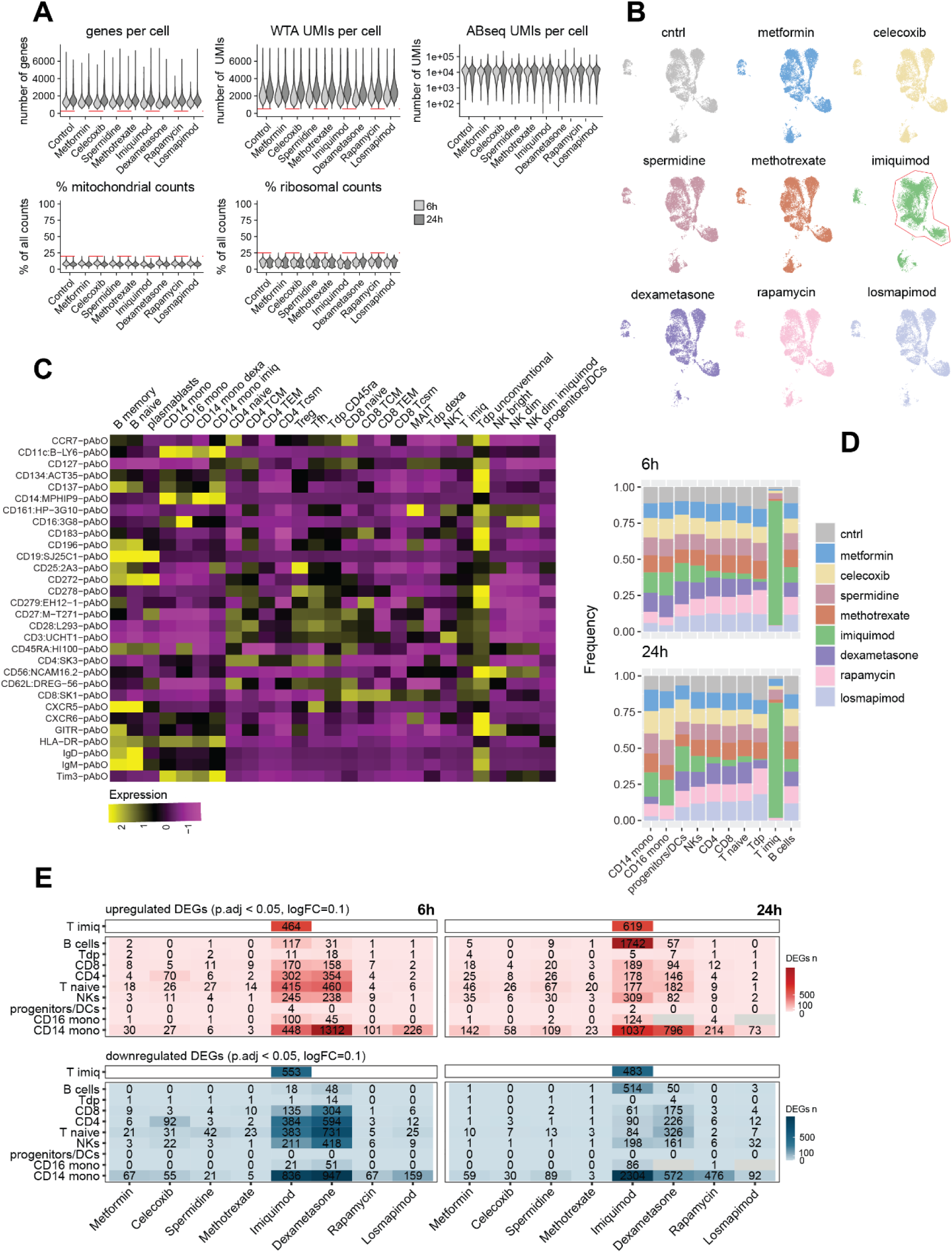
Single-Cell Transcriptomics Reveals Cell-Type-Specific Drug Effects. **A** QC metrics for the generated single-cell multiomics dataset described in Figure 2. **B** UMAP representations of the young human PBMCs single-cell dataset reported in Figure 2 (WTA and ABseq surface proteome integrated profiles) split by drug treatment. **C** Scaled surface proteins expression (ABseq surface proteome) used to annotate cell types and states for the dataset described in Figure 2. **D** Percent contribution of each drug treatment to the final number of cells across the main annotated cell types. **E** Number of DEGs called per each treated versus ctrl comparison across main cell types, drug treatments and time points (p.adj<0.05, logFC_threshold_=0.1). *QC: quality control; WTA: whole transcriptome analysis; UMI: unique molecular identifier; ABseq: surface proteome analysis, DEG: differentially expressed gene*.

**Figure S3-.**
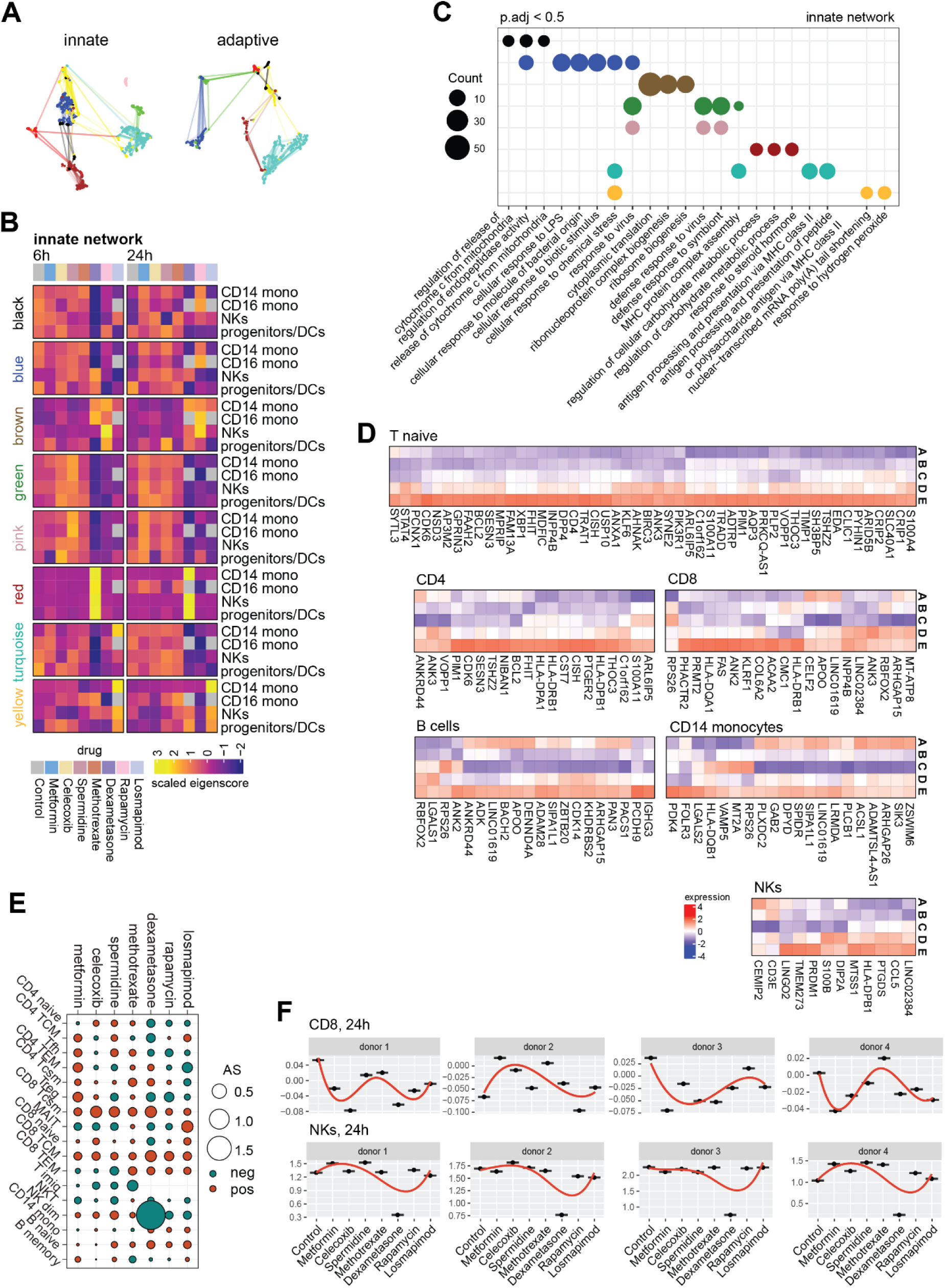
Co-expression Network Analysis Reveals Shared and Distinct Treatment and Drug Effects. **A** hdWGCNA networks generated for innate and adaptive compartments. **B** Heatmap reporting scaled eigenscores across drug treatments and time points for each of the gene co-expression modules identified within the *innate* network. **C** ORA across the gene modules identified within the *innate* network (GO biological process, p.adj<0.05). **D** Heatmap reporting the expression of genes of the cell type-specific *senobase immune aging signatures across age groups in the Terekhova et al. dataset.* **E** Summary ASs describing the anti- or pro-aging effect of different drug treatments on distinct immune cell types and states (granular annotation). **F** Donor-specific ASs dynamics across cell types and states at 24h of treatment in CD8 and NK cells. *ORA: over-representation analysis; GO: gene ontology*.

